# Rainfall-induced microbial resuscitation reveals functional decoupling across biocrust succession

**DOI:** 10.64898/2026.04.20.719569

**Authors:** Raul Roman, Fernando T. Maestre, Estelle Couradeau

**Affiliations:** Department of Ecosystem Science and Management, The Pennsylvania State University, University Park, PA, USA; Instituto Multidisciplinar para el Estudio del Medio Ramon Margalef, Universidad de Alicante, Alicante, Spain; Department of Agronomy, University of Almería, 04120, Almería, Spain; Environmental Science and Engineering, Biological and Environmental Science and Engineering Division, King Abdullah University of Science and Technology, Thuwal, 23955-6900, Kingdom of Saudi Arabia; The Huck Institutes of Life Sciences, The Pennsylvania State University, PA, USA

## Abstract

Dryland ecosystems rely on infrequent rainfall pulses to activate soil microbial communities, yet the fraction and identity of microbes resuscitating after hydration remain unclear. We applied bioorthogonal non-canonical amino acid tagging coupled with fluorescence-activated cell sorting (BONCAT-FACS) and 16S rRNA gene sequencing to identify translationally active bacteria in early (L-BSC) and late (D-BSC) successional cyanobacteria-dominated biocrusts subjected to simulated rainfall under light and dark conditions. Our results reveal that only a small subset of the microbial community resumes activity within six hours, with higher active cell abundances in mature crusts. Microbial activity patterns were largely independent of light exposure and showed partial decoupling from total community composition, indicating that presence does not predict short-term function. These findings suggest that biocrust maturity shapes microbial activation dynamics and that functional responses to precipitation pulses are governed by a conserved pool of fast responders, informing predictions of dryland soil microbiome resilience under changing precipitation regimes.

## 1. Introduction

In drylands, the largest terrestrial biome on Earth, water scarcity constrains biological activity and ecosystem functioning (Collins et al., 2014). In these systems, sporadic precipitation causes brief periods of soil microbial activity, where microbes undergo a pulse regime characterized by long dormancy phases followed by rapid activation upon wetting (Lennon and Jones, 2011; Schimel, 2018). However, it remains difficult to determine both the fraction and the identity of the desert soil microbiome that rapidly resuscitates after quiescence, limiting our ability to link soil functions to microbial activity. With climate models predicting an increased frequency of extreme precipitation events in drylands (IPCC, 2021), understanding these microbial responses is increasingly critical for predicting soil microbiome functioning and biogeochemical processes under climate change scenarios (Maestre et al., 2015).

Desert-inhabiting biological soil crusts (biocrusts) are ideal model systems for exploring coordinated microbial activity after rewetting. These are complex soil surface communities resulting from the close association of photoautotrophs (e.g., cyanobacteria, lichens, or mosses), heterotrophs, and mineral soil particles (Belnap, 2003; Weber et al., 2022). Biocrusts cover approximately 30% of the global dryland area (Rodriguez-Caballero et al., 2018), acting as a mantle of fertility by fixing atmospheric carbon and nitrogen and incorporating these into the soil surface (Chamizo et al., 2012; Barger et al., 2016). Biocrust constituents are well adapted to the pulsed nature of drylands mainly through their capacity to survive prolonged desiccation (Green et al., 2011; Garcia-Pichel, 2023) and quickly transition from quiescence to activity during short hydration events (Xu et al., 2021). This rapid resuscitation has been well characterized in cyanobacteria, which possess several adaptations for revival following hydration and for anticipating forthcoming desiccation (Rajeev et al., 2013; Couradeau et al., 2019a; Xu et al., 2025). In contrast, the identity and fraction of heterotrophic microbiota that quickly become active during these short wetting events remain incompletely resolved, despite growing evidence from transcriptional studies indicating that early resuscitation responses are taxon-specific (Angel and Conrad, 2013; Karaoz et al., 2018; Baubin et al., 2022).

Biocrust development often begins with surface colonization by pioneer, motile filamentous cyanobacteria such as *Microcoleus vaginatus* (Garcia-Pichel et al., 2009), forming light-colored biocrusts (L-BSC). This is typically followed by the establishment of light-adapted sessile heterocystous cyanobacteria such as *Nostoc*, *Scytonema,* or *Tolypothrix* (Belnap et al., 2004), resulting in the formation of dark-pigmented crusts (D-BSC). These successional transitions are associated with increases in microbial biomass, diversity, structural complexity, and photosynthetic and nitrogen-fixation capacities (Belnap et al., 2008; Kuske et al., 2012; Maier et al., 2022). Because cyanobacteria rapidly resume photosynthesis within minutes after wetting under daylight (Rajeev et al., 2013), heterotrophic microbial activation during hydration pulses is often assumed to be closely linked to this activity (Karaoz et al., 2018). Consequently, daylight rainfall events are expected to promote stronger or distinct heterotrophic activation than nocturnal events, during which photosynthesis is absent, and nitrogen fixation is diminished (Belnap et al., 2001). However, it remains unclear whether light exposure during brief rewetting events directly influences microbial activity, or if such coupling is weak or absent in the short term.

Distinguishing active microorganisms from dormant or relic populations is critical for addressing these questions, especially since bulk DNA-based measurements may capture dormant cells, relic DNA, or DNA from non-viable organisms (Carini et al., 2016). Recently, bioorthogonal non-canonical amino acid tagging (BONCAT) has emerged as a powerful approach to probe microbial activity in various environments (Hatzenpichler et al., 2014; Reichart et al., 2020). This method labels translationally active microorganisms by incorporating non-canonical amino acids into newly synthesized proteins, allowing their subsequent recovery and identification via fluorescence-activated cell sorting (FACS) and sequencing (Dieterich et al., 2006; Hatzenpichler and Orphan, 2015). BONCAT-FACS has recently been applied to recover and identify active cells from soils (Couradeau et al., 2019b; Trexler et al., 2023; Harris et al., 2025), providing new opportunities to interrogate microbial resuscitation following hydration in dryland ecosystems and to assess the extent to which microbial presence, as measured by bulk DNA sequencing, reflects short-term translational activity.

Here, we used BONCAT-FACS coupled with 16S rRNA gene sequencing to identify the fraction and identity of microbes that resume translational activity within hours after a simulated rainfall event. We compared early-(L-BSC) and late-successional (D-BSC) cyanobacteria-dominated biocrusts incubated under light (Sun) or darkness (Night) to evaluate whether light exposure during rewetting alters the composition of active communities and whether this response varies across successional stages. We hypothesized that (i) mature crusts host higher total and active microbial abundances, (ii) if heterotrophic activation is linked to photosynthesis, light exposure during rewetting should shift both the fraction and composition of active populations, and (iii) if heterotrophic activation is largely independent of immediate photosynthetic activity, the composition of active populations should remain similar across light conditions and only partially track successional turnover in bulk DNA communities.

## 2. Materials and methods

### 2.1 Site and biocrust sampling description

Biocrust samples were collected at the Green Butte Site (38.720160° N, 109.683221° W), a recreational and cattle grazing area located in Moab (Utah, USA) and situated approximately 1350 meters above sea level. The area is characterized by warm summers and cold winters, with frequent frost and snowfall and an average annual precipitation of 228 mm. Biocrusts are widespread in open spaces and are primarily dominated by cyanobacteria, including filamentous species such as *Microcoleus vaginatus* and *Microcoleus steenstrupii*. *M. vaginatus* is often found in lighter, early successional crusts (L-BSC), whereas *M. steenstrupii* is found in more mature, darker crusts (D-BSC) dominated by heterocystous cyanobacteria such as *Nostoc*, *Scytonema*, and *Tolypothrix* (Couradeau et al., 2016). Fourteen replicates of each biocrust maturity type (n = 28) were collected separately within a 1m x 1m sampling area for each biocrust type. Biocrust samples were collected by pressing circular sterile containers (2.5 cm diameter, 1 cm depth) into the soil and extracting the sample using a sterile spatula underneath the crust. Samples were air-dried upside down, transported to Penn State University (Pennsylvania, USA), and maintained in the dark at 20% relative humidity with drierite desiccant until experimentation.

### 2.2 Biocrust wet-up experiment

Each biocrust microcosm (∼ 4-5 grams) was aseptically unpacked and pre-acclimated for one week under dry conditions in a growth chamber to a 12:12 h light/dark cycle before rainfall simulation. During the light phase, samples were illuminated with a photosynthetically active radiation of 400 µmol photons m□² s□¹. Following acclimation, each microcosm was subjected to a simulated 3 mm rainfall event, receiving 1.36 ml of sterile 1 mM L-homopropargylglycine (HPG) solution (Click Chemistry Tools; Scottsdale, AZ, USA), which corresponds to ∼ 77.5 % of the soil water-holding capacity. HPG is a methionine analog containing an alkyne terminal, allowing downstream click chemistry to fluorescently label newly synthesized proteins (Hatzenpichler and Orphan, 2015). A 3 mm rainfall event was selected to evaluate microbial activation under a low-intensity and frequent precipitation pulse typical of the study site, where more than 61% of the events are equal to or below 3 mm. Half of the microcosms were incubated at 25 °C under light conditions (Sun treatment), while the other half were incubated at the same temperature in complete darkness (Night treatment). Samples were incubated for 6 hours, a time interval expected to capture translational activation of existing ribosomes and to minimize the effect of new biomass generation (Camillone et al., 2026). After 6 hours of incubation, each microcosm was destructively sampled and divided into two subsamples: one for counting viable cells and probing microbial activity through the BONCAT-FACS pipeline, and the other for bulk DNA extraction (Figure 1). At harvest, the subsample undergoing the BONCAT-FACS pipeline was transferred to a 15 mL conical tube containing 5 mL of 0.02% Tween 20 in 1X PBS and vortexed at 3000 rpm for 5 minutes to extract cells. The supernatant containing cell suspensions was stored at −20 °C in 1.25 mL aliquots containing 10% glycerol until further processing. Biocrust subsamples used for retrieving bulk DNA were immediately frozen in liquid nitrogen and stored at −80°C until nucleic acid extraction.

**Figure 1.**
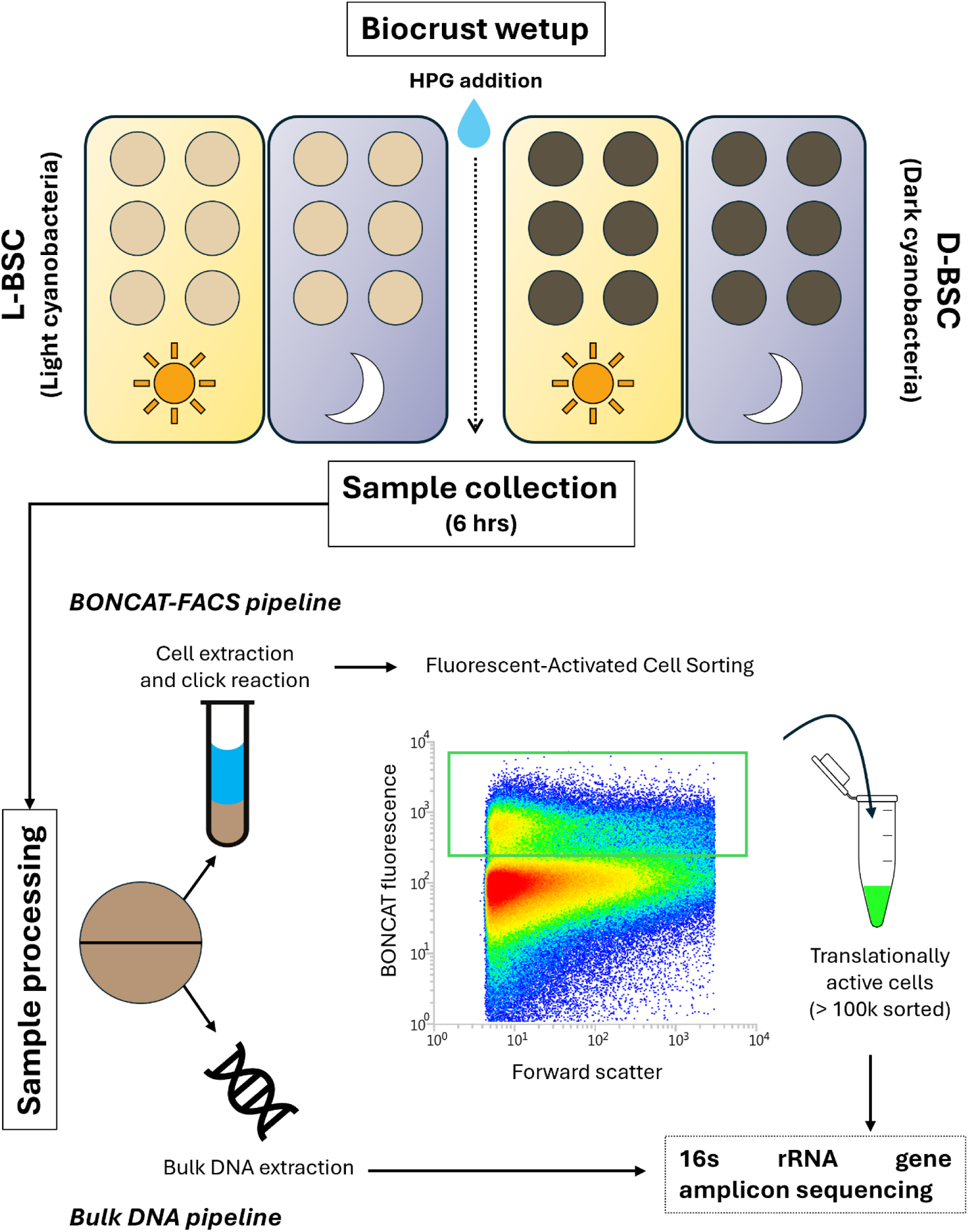
Experimental workflow. Cyanobacteria-dominated biocrusts from early (L-BSC) and late (D-BSC) successional stages received a 3 mm simulated rainfall event containing HPG (total n = 28). Half of the samples were incubated under light (Sun treatment), whereas the other half were incubated in dark conditions (Night treatment). After 6 hours, samples were collected and divided into two sections: one underwent the BONCAT-FACS pipeline, which included cell extraction in saline buffer, HPG click reaction, and fluorescent-activated cell sorting, and the other part was used for bulk DNA extraction. Both fractions were analyzed by 16S rRNA gene amplicon sequencing (see below) to determine the composition of Bulk DNA and BONCAT-active bacterial communities.

### 2.3 Probing microbial activity with BONCAT

BONCAT was used to track cellular activity within the biocrust microbiome by incorporating HPG into newly synthesized proteins. A subsample aliquot was thawed at 4°C for 1 hour and processed following a modified click-chemistry protocol adapted from Reichart et al. (2020). Samples were gently vortexed and centrifuged at 500 × g for 5 min to pellet soil particles, after which the supernatant was recovered and passed through a 35 μm cell strainer to remove residual debris. Cells were then pelleted by centrifugation at 14,000 × g for 5 min and resuspended in 300 μL of 1x PBS, followed by a brief vortex to ensure thorough mixing. Subsequently, a copper-catalyzed click reaction was conducted to bind the HPG to a fluorescent probe (FAM picolyl azide dye, Ex/Em 490/510 nm), making all translationally active cells fluorescent, as described by Couradeau et al (2019). Briefly, 200 μL of the reaction mix was added to each sample and incubated for 30 min at room temperature in the dark. The reaction mix consisted of 5 μl CuSO4 (100 μM final concentration), 10 μl tris-hydroxypropyltriazolylmethylamine (THPTA, 500 μM final concentration), and 3.3 μl FAM picolyl azide dye (5 μM final concentration), buffered with 50 μl freshly prepared 5 mM sodium ascorbate and 50 μl 5 mM aminoguanidine HCl in 1x PBS, and brought to 880 μl with PBS. All reagents were purchased from Click Chemistry Tools (Arizona, USA). Following incubation, excess dye was washed off by centrifugation at 14,000 x g for 5 min. Pellets were washed three times with 1 mL of 1X PBS and finally resuspended in 500 μL PBS for downstream processing.

### 2.4 Flow cytometry and sorting of translationally active cells

The proportion of translationally active cells (BONCAT-active) relative to the viable microbial community was quantified using flow cytometry on a BD LSRFortessa Cell Analyzer (Becton, Dickinson and Company). To distinguish viable cells from soil particles, broken cells, debris, and free DNA, samples were stained with SYTO59 DNA counterstain (0.5 μM; Invitrogen; Ex/Em 622/645 nm) for 5 min in the dark. Instrument thresholds were set to detect particles larger than 0.2 μm using the Flow Cytometry Sub-Micron Particle Size Reference Kit Beads (Invitrogen). The instrument was set up to capture the BONCAT fluorescence via the FAM picolyl azide dye (Ex/Em 490/510 nm) on the green channel, using a 488 nm blue laser, and SYTO59 fluorescence in the red channel using a 630 nm red laser. Viable cells were identified by gating SYTO59-positive events against unstained control. BONCAT-active cells were then enumerated from within the SYTO59-positive population using a nested gate on the green channel. A water-incubated and HPG-free control sample was used to define background fluorescence associated with the click reaction on the green channel, and the false-positive recovery was set at 0.2%. The same gating strategy was subsequently applied to collect BONCAT-active cells using FACS on a MoFlo Astrios Cell Sorter (Beckman Coulter). BONCAT-active cells were sorted from each sample in 1.5 mL Eppendorf tubes, and approximately 100,000 active cells were collected per sample for DNA extraction. Total cell density was independently estimated by counting SYBR Green-positive events per microliter, using unstained samples as a control. This BONCAT-FACS pipeline enabled the quantification of the proportion of BONCAT-active cells relative to viable cells (Harris et al., 2025), as well as estimates of total and active microbial cell abundances per square meter for each biocrust type and treatment condition (night vs. sun).

### 2.5 DNA extractions and 16S rRNA gene amplicon sequencing

We extracted bulk total DNA from 250 mg of dry biocrust sample with the DNeasy PowerSoil Pro Kit (Qiagen). To enhance DNA collection, samples were added to bead tubes and disrupted at 30 Hz for 1 minute in a Tissuelyser II using a tube adaptor (QIAGEN, Aarhus, Denmark). Our BONCAT-active library preparation was adapted from Reichart et al. 2020. Sorted cells were centrifuged at 14000□×□g for 10□min at 4□°C. Subsequently, the supernatant was removed by gentle aspiration. The pelleted cells were lysed using PrepGEM (zyGEM, Charlottesville, VA, USA) chemical lysis in 5□µl reactions, following the manufacturer’s instructions. Each tube received 0.5□µL of 10X Green buffer, 0.084□µL of PrepGEM, 0.084□µL of lysozyme, and 4.35□µL of water. Subsequently, tubes were vortexed for 1 min to detach the cells from the tube walls, and pulse centrifuged at 500 x g. The 5 µL mix was transferred to 0.2 µL PCR tube strips and incubated in a thermocycler at 37□°C for 30□min, followed by 75□°C for 40□min.

The 515f-Y (GTGYCAGCMGCCGCGGTAA, Parada 2016) and 806R-B (GGACTACNVGGGTWTCTAAT, April 2015) primers were used to amplify the 16S rRNA gene from BONCAT-active cell lysate and bulk total DNA. PCR reactions occurred in a final volume of 25 µL, including 12.5□µl of the 2X Platinum™ SuperFi™ PCR Master Mix, 1.25□µl of the forward primer (at 10□µM), 1.25□µl of the reverse primer (at 10□µM), 5 µl of 5X SuperFi™ GC Enhancer, and 5□µl of cell lysate or DNA template. The PCR conditions involved an initial denaturation step at 98□°C for 2□min, followed by 30 cycles consisting of a 10□s denaturation step at 98□°C, a 20□s annealing step at 55.4□°C, a 15□s elongation step at 72□°C, and a final elongation step of 5□min at 72□°C. Library preparation and indexing were conducted at The Huck Life Science Genomics Core. Libraries were sequenced on an Illumina MiSeq platform using 250 bp x 250 bp paired-end sequencing with 15% PhiX.

### 2.5 16S rRNA gene amplicon-sequencing data processing

Demultiplexed FASTQ files were processed with Cutadapt (v1.18) to remove primers. Primers were removed in paired-end mode using anchored 5’ adapters for the primers at the read starts and non-internal 3’ adapters for the reverse complements at the read ends. Trimmed reads were processed in R using DADA2 (v1.26, Callahan et al., 2016). After quality profile inspection, forward and reverse were truncated at 230 and 200 base pairs (bp), respectively. After truncation, reads with more than two (forward) or five expected errors (reverse) were discarded (maxEE = c(2,5)), and reads with ambiguous bases were removed. After sequence inference and chimera removal (method = consensus), we used the DADA2 native naïve Bayesian classifier method to assign taxonomy, pre-trained on the SILVA database (v138.2_toGenus). To reduce redundancy at the ASV level, ASV sequences were oriented and aligned in DECIPHER and clustered at 97% identity using hierarchical clustering (TreeLine, UPGMA, cutoff = 0.03), selecting the most abundant ASV per cluster as the representative sequence. ASVs belonging to chloroplasts, or mitochondria were removed. A subset containing Cyanobacteriota was reserved for the exploration of relative abundance between biocrust types and fractions (Figure S1). Due to the known limitations of flow cytometry for capturing large filamentous organisms (Trexler et al., 2023) and to focus exclusively on non-photoautotrophic bacteria, we filtered out Cyanobacteriota for the rest of the bioinformatic analyses. This quality-controlled dataset was then subsequently filtered to remove extra rare taxa by keeping only those that: i) were present in at least 3 samples, and ii) had at least 50 reads.

### 2.6 Statistics of microbial community composition

We conducted the statistical analysis in R (v. 4.5.1). The filtered dataset was used to test differences in microbial composition across fractions (Bulk DNA vs BONCAT-active), biocrust type, and incubation conditions. Bray-Curtis dissimilarities between samples were calculated from relative abundance tables using the vegdist() function of the R package vegan (Oksanen et al., 2026) and visualized using a non-metric multidimensional scaling (NMDS) ordination that included 95% confidence ellipses drawn around fraction groups. The effects of individual factors and their interactions on community composition were tested using a PERMANOVA via the function adonis2() from the same package. β-dispersion differences were quantified using the Euclidean distance of each sample to its group centroid using vegan::betadisper() and vegan::permutest() to test for significance among groups. Distances were extracted from the results output and fit to a linear model to evaluate each factor individually, as well as all pairwise interactions among them. Estimated marginal means (EMMs) were computed from that model using the emmeans R package (ref), and pairwise differences among EMMs were tested using Tukey HSD post-hoc comparisons. The relative abundance of bacterial phyla was depicted using a stacked bar plot using the ggplot2 v.3.5.1 package.

Because light exposure had a non-significant influence on microbial composition, data from both incubation conditions were pooled to identify taxa significantly enriched or depleted in the BONCAT-active fraction relative to the Bulk fraction. Differential abundance analyses were conducted using ANCOM-BC2 (v2.2.1, Lin et al., 2024) implemented in R. Independent ANCOM-BC2 models were run at the ASV level for (i) the BONCAT-active vs. Bulk DNA contrast within each biocrust type level and (ii) the L-BSC – D-BSC contrast within each fraction level. Models included a single fixed effect (Fraction or Biocrust type) and were fitted using FDR adjustment (q < 0.05), pseudo_sens = T, struc_zero = F, and the rest of the parameters set to default. The resulting significant log-fold-changes (LFC) estimates were extracted, and the top 20 ASVs with the largest absolute LFC values (10 enriched and 10 depleted per contrast and level) were displayed in bar plots using ggplot2. To evaluate the consistency of microbial responses across contrasts, LFC values were compared between the two levels of each contrast using scatterplots, including those ASVs that were significantly enriched or depleted in one or both levels of the contrast. Spearman correlations were calculated exclusively for the ASVs that were significant in both levels of each contrast to quantify concordance in the magnitude and direction of the enrichment or depletion.

## 3. Results

### 3.1. Differences in microbial abundance and activity across biocrust types and incubation conditions

The abundance of total and active cells differed significantly between biocrust types and incubation conditions (Figure 2). Total microbial abundance was consistently higher in D-BSC compared to L-BSC under both incubation conditions, reaching up to 3-fold higher values in D-BSC incubated under Sun. Within L-BSC, the Night incubation exhibited higher cell abundances than the Sun incubation, whereas no significant differences between incubation conditions were found in the D-BSC (Table S1). For BONCAT-active cells, the biocrust type had a strong effect (F = 34.27, *P* < 0.001), while neither incubation conditions nor the interaction had significant effects (*P* = 0.125 and *P* = 0.545, respectively). Active cell abundances were higher in D-BSC than in L-BSC, showing a proportion of BONCAT-active cells around 7.1-7.6 % of the total community in D-BSC, compared to approximately 4% in L-BSC. Active cell abundances tended to be higher under Night than Sun incubation in L-BSC, but this pattern was absent in D-BSC.

**Figure 2.**
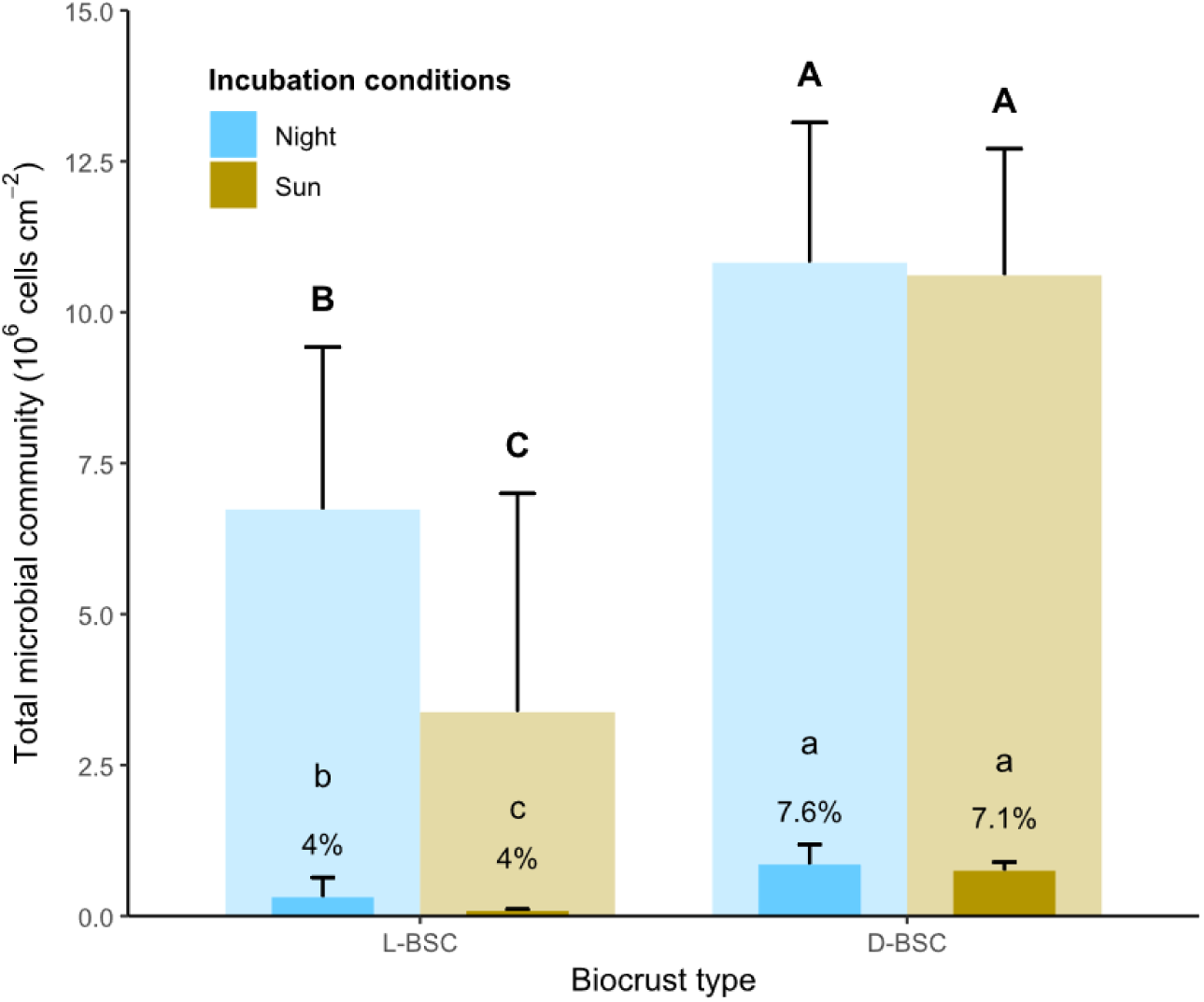
Mean abundance of total and BONCAT-active microbial cells in both biocrust types. Bars represent the total microbial community (semi-transparent) with overlaid BONCAT-active fractions (solid bars). Percentages above error bars correspond to the mean proportion of BONCAT-active cells relative to the total community. Capital and lowercase letters indicate significant differences (ANOVA, *P* < 0.05) among incubation conditions and biocrust types for total and active fractions, respectively. All test results are in Tables S1 and S2.

### 3.2. Fraction and biocrust type as main drivers of microbial community structure

We used BONCAT-FACS combined with 16S rRNA gene sequencing to identify the active members of the microbial community in L-BSC and D-BSC. Non-metric multidimensional scaling (NMDS) based on Bray-Curtis dissimilarity showed a clear separation between BONCAT-active and Bulk DNA fractions, particularly in D-BSC (Figure 3A). Biocrust type also influenced community composition, but the separation between biocrust types was more pronounced in the Bulk DNA fraction. Samples incubated under Sun and Night clustered similarly, indicating that incubation conditions had no significant effect on community composition. PERMANOVA results supported these observations, showing that fraction and biocrust type significantly shaped bacterial composition, explaining 25.4% and 15.8% of the variation, respectively (Table S1). In contrast, incubation condition was not significant, accounting for only about 0.9% of the variation. Additionally, a significant interaction between fraction and biocrust type indicated that the effect of biocrust type on community composition differed between BONCAT-active and total community DNA fractions, consistent with the NMDS ordination patterns. Incubation treatment did not significantly affect the within-group variation (β-dispersion) of biocrust microbiomes (ANOVA, *F* =□0.5169, *P*□=□0.47). β-Dispersion was equal between L-BSC and D-BSC in the bulk DNA fraction, while it differed markedly in the BONCAT-active fraction, with L-BSC showing significantly higher β-dispersion than D-BSC (Figure 3B).

**Figure 3.**
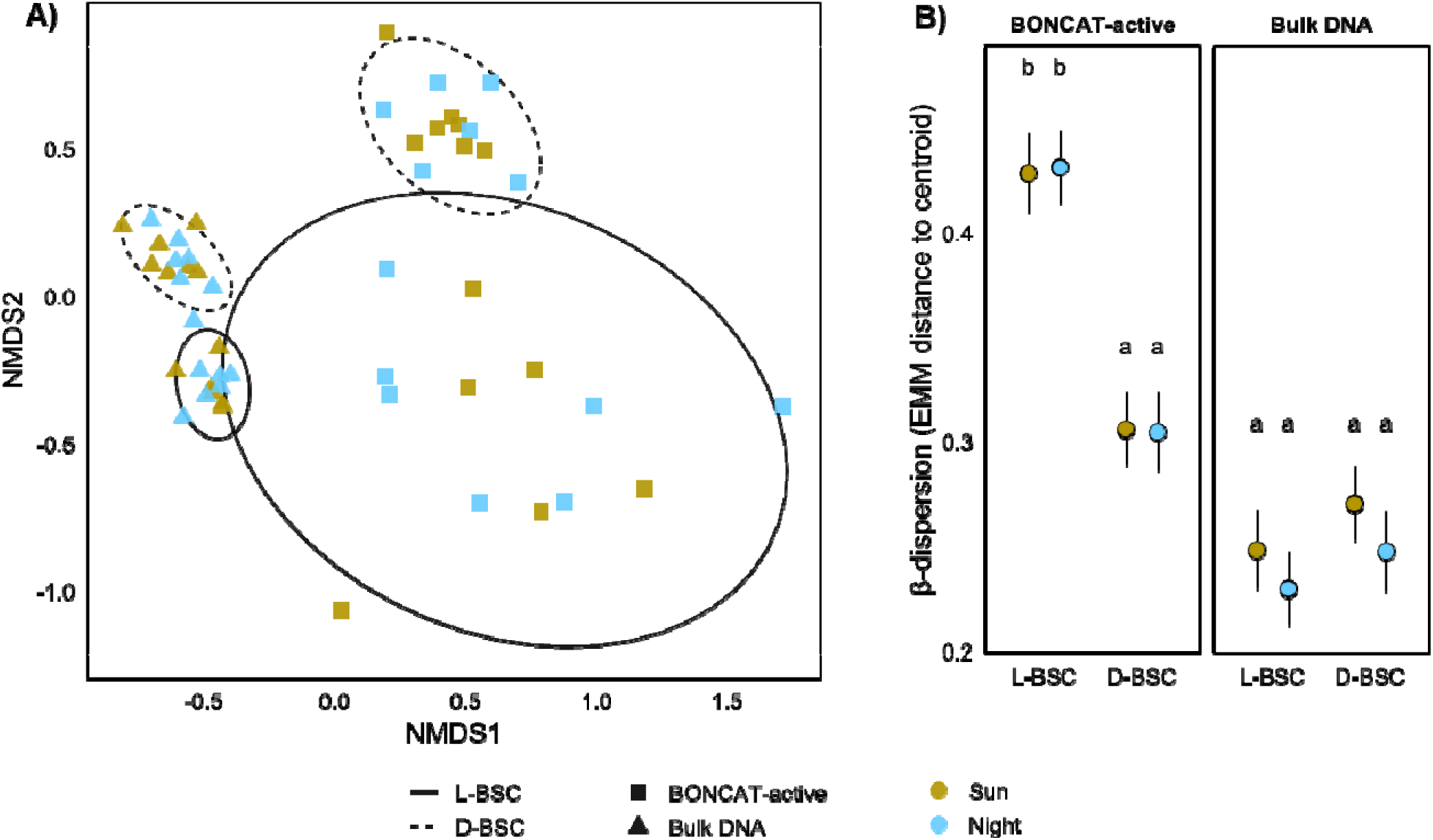
(A) Non-metric multidimensional scaling (NMDS) ordination based on Bray–Curtis dissimilarities showing bacterial community composition by incubation condition (Sun vs Night), with point shapes indicating fraction (BONCAT-active vs Bulk DNA). The 95% confidence ellipses represent biocrust type (L-BSC: solid; D-BSC: dashed). Cyanobacteria were excluded from the ordination due to low representation in the BONCAT-active fraction (Figure S1) and to reduce bias associated with large filamentous organisms (see method section 2.5). (B) Estimated marginal means (EMMs) of β-dispersion, calculated as the Bray-Curtis distance of each sample to its group centroid, error bars represent ±SE, and letters indicate pairwise group differences based on the Tukey HSD post-hoc test.

### 3.3. Differentially abundant taxa between BONCAT-active and Bulk DNA fractions

As light exposure had no significant influence on microbial structure, both incubations were pooled together to analyze the microbial composition between fractions. The relative abundance of the most abundant phyla differed significantly between fractions in both biocrust types. In the Bulk DNA fraction, Pseudomonadota, Actinomycetota and Chloroflexota were the most abundant in both biocrust types, accounting for over 60% of all reads (Figure S2 and S3), suggesting that many microbial cells assigned to these phyla were either dormant or not viable. Conversely, Armatimonadota was more abundant in the BONCAT-active fraction in both biocrusts, whereas Abditibacteriota was significantly higher only in L-BSC.

To determine which taxa drove these compositional differences, we applied ANCOM-BC2 to identify differentially abundant ASVs between fractions (q < 0.05). Across both biocrusts, 488 ASVs significantly differed between fractions, including 102 in the BONCAT-active and 386 enriched in the Bulk DNA fraction (full list in Table S3). ASVs enriched in the Bulk DNA fraction were dominated by Actinomycetota, which accounted for approximately 68 % (263 of 386) of all ASVs in the Bulk DNA fraction. Within this phylum, enrichment was concentrated in a few genera, including *Rubrobacter*, *Nocardioides*, *Pseudonocardia*, *Propionibacterium*, *Geodermatophilus*, and *Blastococcus* (Figure 4, negative LFC). Additional ASVs enriched in this fraction belonged to Pseudomonadota (15 %; 59 ASVs), Chloroflexota (6 %; 23 ASVs), and Bacteroidota (5 %; 20 ASVs). Conversely, most BONCAT-active ASVs belonged to Abditibacteriota, Bacteroidota, and Deinococcota in both biocrust types. Within these, activity was concentrated in a small number of genera, including *Abditibacterium*, *Segetibacter*, and *Deinococcus*, which contained multiple ASVs strongly enriched in the BONCAT-active fraction of both biocrust types, and minor enrichments by *Sphingomonas* and *Solirubrobacter* (Figure 4, Table S3).

**Figure 4.**
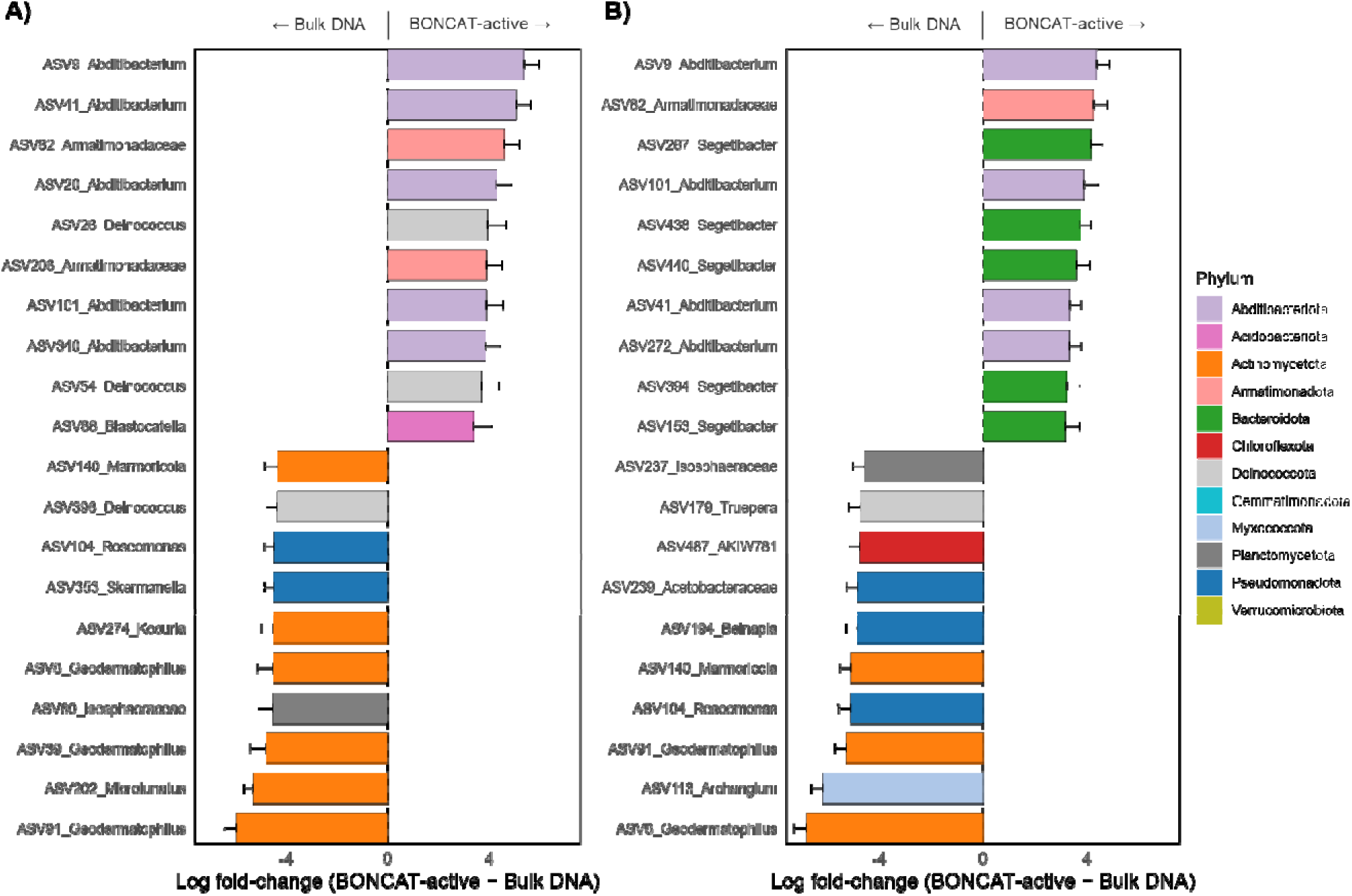
Top 10 differentially enriched ASVs in BONCAT-active versus bulk DNA fractions across biocrust types: A) Light cyanobacteria-dominated (L-BSC) and B) Dark cyanobacteria-dominated (D-BSC). ASVs were selected by the largest absolute LFC for the BONCAT-active versus bulk DNA contrast. Positive LFC values indicate enrichment in the BONCAT-active, whereas negative values indicate enrichment in the Bulk DNA fraction. Only ASVs with significant differential abundance after FDR Benjamini–Hochberg correction (q < 0.05) are shown.

### 3.4. Concordance between translational activity and community composition

To evaluate whether the taxa enriched in different fractions were consistent across biocrust developmental stages, we compared the LFCs estimates for the BONCAT – Bulk DNA contrasts between L-BSC and D-BSC crusts (Figure 5A). Approximately one third of the ASVs showed concordant direction and magnitude of change in both biocrusts, clustering along the 1:1 diagonal and indicating that the same taxa tended to be either translationally active (positive LFC) or present in the bulk community (negative LFC) regardless of biocrust maturity (Spearman ρ = 0.65, p < 0.001). However, the specific composition of active taxa shifted along the successional gradient, since only 21.4% (18 of 84) of ASVs enriched in the BONCAT-active were shared between L-BSC and D-BSC. In contrast, the proportion of ASVs showing shared enrichment in the Bulk DNA fraction was significantly higher (36.4%).

**Figure 5.**
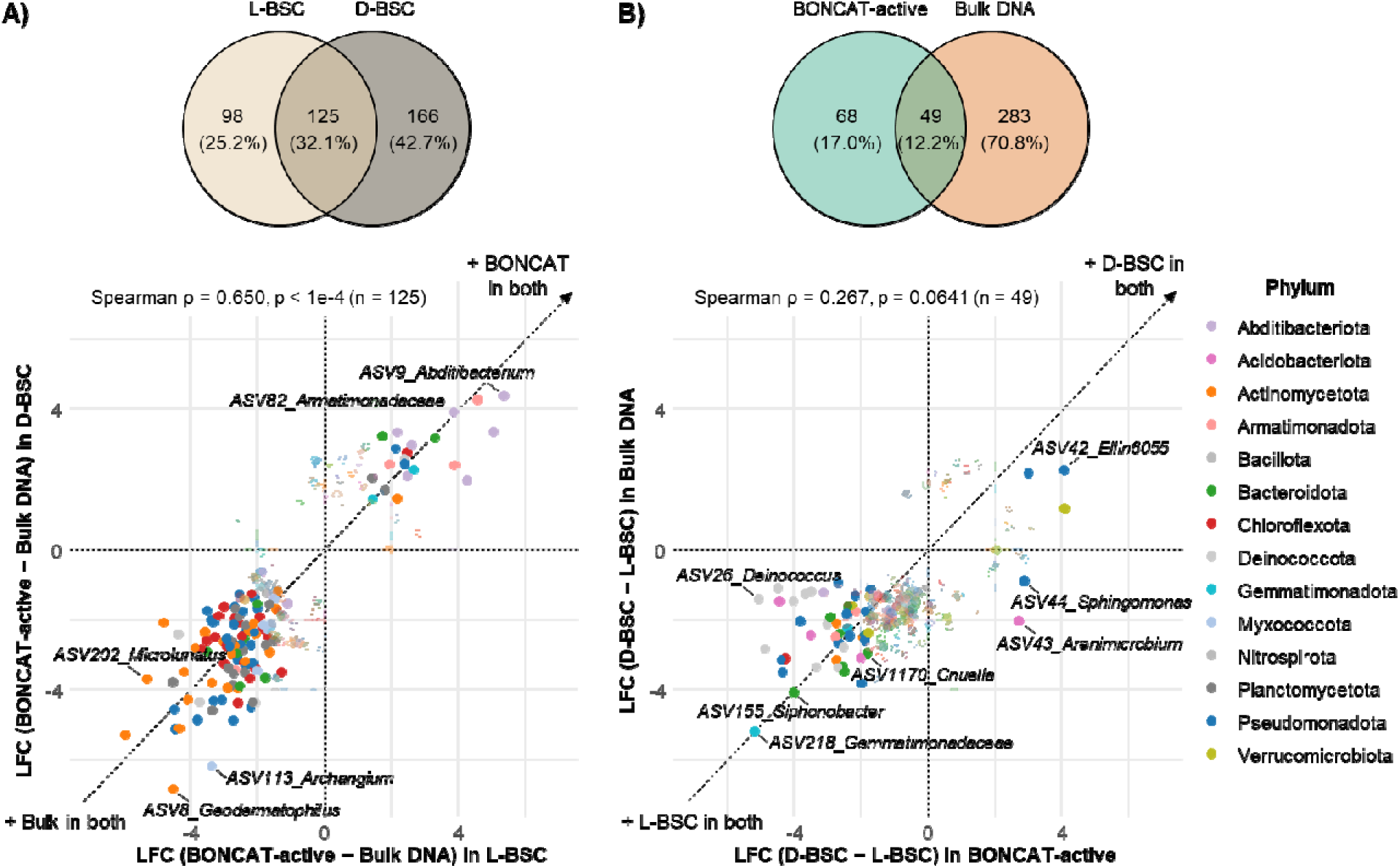
Concordance of differential enrichment patterns across biocrust maturity and activity fractions. Scatterplots compare ANCOM-BC2 log fold-change (LFC) estimates for individual ASVs (each point) between pairs of contrasts. (A) Comparison of the BONCAT-active vs bulk DNA contrast across successional stages: the x-axis shows LFCs in L-BSC, and the y-axis show LFCs in D-BSC for the same ASVs. Positive LFC values indicate ASVs enriched in the BONCAT-active fraction, whereas negative values indicate enrichment in the bulk DNA fraction. (B) Comparison of the D-BSC vs L-BSC contrast between fractions: the x-axis shows LFCs estimated from the BONCAT-active fraction, and the y-axis shows LFCs from bulk DNA. Positive LFC values indicate ASVs enriched in D-BSC (mature) relative to L-BSC (early), whereas negative values indicate enrichment in L-BSC. In both panels, ASVs significant in at least one of the two compared conditions (q < 0.05) are shown; light-colored points are significant in only one condition, and dark-colored points are significant in both. The horizontal and vertical zero lines separate enrichment directions, and quadrants summarize concordant (top-right, bottom-left) versus divergent (top-left, bottom-right) responses. Venn diagrams show the overlap in significant ASVs between the compared conditions. Spearman correlations (ρ) quantify agreement in enrichment magnitude and direction using only ASVs significant in both conditions.

To test whether compositional changes measured in the Bulk DNA community along the successional gradient produced parallel responses in translational activity, we compared D-BSC – L-BSC contrast between BONCAT-active and Bulk DNA fractions (Figure 5B). Only a small group of ASVs (12.2%) showed concordant responses between fractions, and the relationship between them was weak and not significant (Spearman ρ = 0.27, p = 0.064), revealing that total community shifts linked to biocrust maturity were not reflected in equivalent changes in active taxa. Nevertheless, a few distinct outliers showed contrasting response types. For example, members of the genus *Ellin6055* showed positive LFCs in both fractions, suggesting an increase in both activity and abundance with maturity, whereas *Cnuella*, *Massilia*, *Noviherbaspirillum* and members of Gemmatimonadaceae increased presence and activity in early-successional biocrusts (negative LFCs). However, the scattered distribution of points and low fraction concordance confirm that most taxa vary independently in abundance and activity, supporting the idea that bulk microbial presence and short-term metabolic activity signatures are partially decoupled across the biocrust successional gradient.

## 4. Discussion

### Microbial abundance and activity are controlled by biocrust maturity

Consistent with our first hypothesis, both total active and cell abundances differed between biocrust developmental stages. D-BSC exhibited higher cell abundances than L-BSC, consistent with previous findings that report increased cell abundance with greater photoautotrophic biomass (Maier et al., 2018). Our study builds on previous research and demonstrates for the first time that biocrust maturity not only increases microbial abundance but also hosts larger populations of translationally active members. Increased microbial activity may also explain the higher respiration rates consistently reported in more developed biocrusts (Grote et al., 2010; Maier et al., 2018; Lopez-Canfin et al., 2022). These functional differences are typically attributed to greater organic carbon and microbial biomass. However, our results suggest that differences in the size of the active fraction may also play a role, given that BONCAT-active cell counts are positively correlated with respiration rates in soils (Camillone et al., 2026).

Despite these differences, only a small fraction of the community (4 – 7%) resumed translational activity within the 6-hour pulse (Figure 2). Nevertheless, these results are well above the proportions reported in bulk soil and rhizosphere using the same methodology (Harris et al., 2025), and are within the range of active cell numbers reported for other soils using different methods (Blagodatskaya et al., 2013). Recent studies combining single-cell ²H_2_O labeling and NanoSIMS imaging suggest a much faster and more extensive resuscitation of the heterotrophic community in biocrusts, with over 75% of cells showing anabolic activity after the same time interval (Imminger et al., 2024). While these findings indicate that most cells resume minimal repair or maintenance activity after hydration, our results show that only a small subset engages in detectable translational activity during brief wetting. Because BONCAT-FACS detects translational activity of cells not necessarily undergoing division, our approach is better suited to capture activation dynamics that otherwise might not be detected by growth-based methods. Looking forward, this methodology can be applied to identify rare and active species that may disproportionately contribute to short-term metabolic fluxes in biocrusts (Friedline et al., 2025), an outstanding research question in the field that has remains unanswered.

### Short-term microbial activation is decoupled from real-time photosynthesis activity

Contrary to our second hypothesis, light exposure had no significant effect on microbial composition and only minor effects on microbial activity. Incubation under Sun or Night (no photosynthesis) conditions in D-BSC resulted in only minor, non-significant variations in activity (Figure 2), with active cell proportions consistently around 7%. In L-BSC, light exposure led to lower cell abundances, affecting the total number of active cells, but the proportion of active cells remained stable (∼4%) across incubation conditions. Similarly, light exposure had a minimal influence on microbial community composition (<1% observed variance), consistent with the observed cell activity patterns and indicating that sunlight during short-term wetting has little impact on which taxa reactivate. These results indicate that microbial cell abundance and activity following a single wet-up event are decoupled from cyanobacterial photosynthetic activity, contrary to our initial expectation.

This asynchronous response is somewhat unexpected and challenges the prevailing view of interdependent resuscitation between photoautotrophs and heterotrophs in cyanobacteria-dominated biocrusts (Baran et al., 2015; Swenson et al., 2018). However, we cannot rule out the possibility of light-dependent coupling in the uppermost biocrust layer, the photic zone (0–1 mm depth), which is an O_2_ supersaturated environment (Garcia-Pichel et al., 2023). In this zone, bundle-forming cyanobacteria actively attract copiotrophs and diazotrophs, facilitating carbon-for-nitrogen mutualistic exchanges that could enable coupled activity (Couradeau et al., 2019a; Nelson et al., 2022). Further research is needed to elucidate these microscale metabolic interactions, for example, by using BONCAT to identify translationally active members of the cyanosphere and track proteomic changes across diel cycles (Loynd et al., 2026).

### BONCAT-FACS reveals early responding taxa across biocrust succession

The clear separation between fractions observed in the NMDS (Figure 3A) reveals a strong distinction between presence and activity in both biocrust types. The large compositional differences observed between L-BSC and D-BSC in the Bulk DNA fraction were expected and align with prior work showing clear taxonomic changes along cyanobacteria-dominated maturity gradients (Couradeau et al., 2016; Roncero-Ramos et al., 2020). Notably, the significant interaction between fraction and biocrust type (Table 1) reveals that biocrust maturity modulates the relationship between bulk presence and activity—a new finding. As crusts mature, their active communities diverge more from their total DNA pool (Figure 2). This separation may be attributable to increasing cyanobacterial diversity and associated heterotrophic community complexity (Maier et al., 2018), stronger vertical stratification, and increased accumulation of dormant cells or relic DNA over time in mature crusts, a common phenomenon in soils (Carini et al., 2017). Additionally, β-dispersion of BONCAT-active populations was markedly lower in D-BSC than in L-BSC, suggesting that mature communities tend to converge in their composition of active bacteria, whereas L-BSC exhibits a more heterogeneous composition (Xu et al., 2020).

**Table 1.**
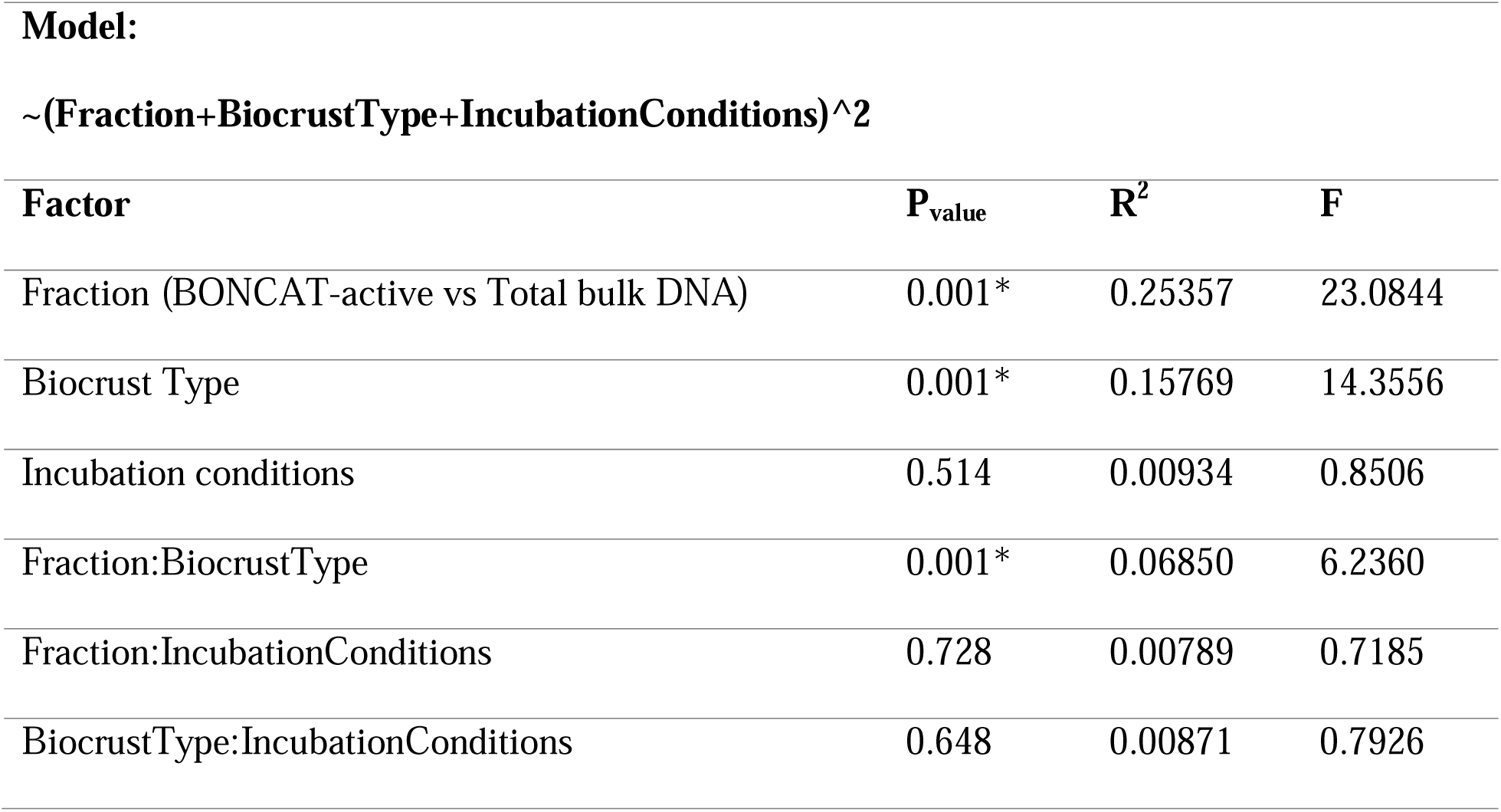
PERMANOVA results based on Bray-Curtis dissimilarities testing the effects of microbial fraction, biocrust type, incubation conditions, and their interaction on microbial community composition. Asterisks (*) indicate p-values where factors significantly impacted microbial community composition, while R^2^ values reflect the percentage of variation that can be explained due to the experimental factor.

**~(Fraction+BiocrustType+IncubationConditions)^2**
| **Factor** | **P<sub>value</sub>** | **R<sup>2</sup>** | **F** |
| --- | --- | --- | --- |
| Fraction (BONCAT-active vs Total bulk DNA) | 0.001* | 0.25357 | 23.0844 |
| Biocrust Type | 0.001* | 0.15769 | 14.3556 |
| Incubation conditions | 0.514 | 0.00934 | 0.8506 |
| Fraction:BiocrustType | 0.001* | 0.06850 | 6.2360 |
| Fraction:IncubationConditions | 0.728 | 0.00789 | 0.7185 |
| BiocrustType:IncubationConditions | 0.648 | 0.00871 | 0.7926 |

Our analyses found that Bulk-enriched ASVs mostly belonged to Actinomycetota, which have been previously found in many other biocrust communities (Angel and Conrad, 2013; Van Goethem et al., 2019; Baubin et al., 2022). Whereas a few ASVs belonging to Actinomycetota were also enriched in the BONCAT-active fraction, their much higher abundance in the Bulk fraction of both crusts suggests that many members of this phylum (e.g., *Geodermatophilus, Nocardioides, Pseudonocardia*) exhibit a slow response to wetting or remain dormant during short rainfall pulses, as previously reported (Trexler et al., 2023). This group includes well-known oligotrophic bacterial genera ubiquitous to cyanobacteria-dominated biocrusts (Miralles et al., 2020; Zhou et al., 2024) but are typically absent from the cyanosphere (Couradeau et al., 2019a; Nelson et al., 2021), which may partially explain their reduced activity during short-term hydration. Actinomycetota also exhibits physiological adaptations to desiccation, such as sporulation (Setlow, 2007), which allows for extremely long dormant periods at the cost of a very long pulse to enable germination (Dworkin and Shah, 2010). In contrast, the BONCAT-active fraction was predominantly enriched by Abditibacteriota, Bacteroidota, and Deinococcota in both biocrusts. Remarkably, *Abditibacterium*, the first cultivated representative of the candidate phylum FBP isolated from Antarctic mineral soils (Tahon et al., 2018), showed a strong enrichment in the BONCAT-active of both L-BSC and D-BSC. To our knowledge, this work represents the first evidence of *Abditibacterium* as an active member of the biocrust community. Other active genera, including *Cnuella*, *Massilia*, or *Noviherbaspirillum*, were preferentially enriched in L-BSC (Table S3). The same taxa have been found to respond to short rainfall pulses (5 hr) in very distant geographical early-successional biocrusts from the Chihuahuan Desert (Kut and Garcia-Pichel, 2024), suggesting that they have specific adaptations to preserve cell structure after recurrent hydration-desiccation and preparedness for resuscitation (Imminger et al., 2024).

Our observations are also consistent with a general phylogenetic organization of nimble vs torpid responders in biocrusts postulated in the expanded pulse-reserved paradigm of Garcia-Pichel and Sala (2022). They also align with recent evidence showing that members of Actinomycetota were mainly associated with long-pulse responder strategies, while *Cnuella*, *Massilia*, or *Noviherbaspirillum* were associated with the short-pulse strategies (Kut and Garcia-Pichel, 2024). Our BONCAT-FACS builds on this framework by directly identifying taxa that rapidly resume protein synthesis following short hydration pulses. Now that BONCAT-live permits the isolation and cultivation of BONCAT-active sorted cells (Mulay et al., 2025), future research could focus on culturing early and late responders to further confirm these differences in life strategy.

### Activity patterns are consistent across biocrust maturity but decoupled from total community structure

Beyond taxa-specific activation patterns, a substantial portion (>20%) of the population significantly enriched in the BONCAT-active fraction was shared by both early and mature biocrusts, with most active ASVs clustering along the 1:1 diagonal in both crusts (Figure 5A). These results suggest that biocrusts share a pool of fast responders that reactivate metabolism upon wetting, regardless of their taxonomic composition, likely reflecting a form of conserved functional redundancy typical of dryland microbiomes (Sauma-Sánchez et al., 2024). However, D-BSC crusts still exhibited a larger and more diverse population enriched in their active fraction (Table S3), mainly dominated by Actinomycetota, Bacteroidota, and Pseudomonadota, consistent with higher niche specialization and increased functional specificity and complexity (Jiang et al., 2025).

As we hypothesized, we found a weak correlation between presence and activity across biocrust development (Figure 5B, Spearman ρ = 0.27, *P* = 0.096), indicating a functional decoupling between microbial abundance and activity across the maturity gradient. This decoupling demonstrates that dominant or persistent taxa in the bulk community, as determined by traditional 16S rRNA methods, are not necessarily responding to fast rehydration. Because the most frequent precipitation events in the study area fall in the 0.25–2.87 mm range, which results in hydration periods equal to or lower than our experiment wet duration, we argue that this short-term metabolic response is realistic and captures the microbial resuscitation process typical of cyanobacteria-dominated biocrusts. Considering the most likely predicted shift towards more frequent but smaller summer precipitation events in the southwestern US (Weltzin et al., 2003), we anticipate these groups will be favored, with unknown consequences for biocrust-mediated ecological functions in desert ecosystems.

## 5. Conclusion

Our results demonstrate that only a small fraction of the cyanobacteria-dominated biocrust microbiome resumed translational activity following a single wet-up event. Estimated cell abundances, microbial activation, and community structure were strongly influenced by biocrust maturity, whereas light availability had no significant effect. Although early and mature biocrusts shared a pool of active taxa, early-successional crusts exhibited higher β-dispersion, whereas more mature biocrusts converged toward a more consistent and diverse active microbial assemblage. We further demonstrate that the dominant taxa retrieved in the bulk community and active microbes signatures are partially decoupled, indicating that presence alone is not a good predictor of short-term functional responses. Consequently, metabolic fluxes following short rainfall events in desert soils are likely driven by a distinct subset of fast-responding taxa, rather than by the dominant taxa identified in bulk surveys. Under increased human pressure and changing precipitation regimes, regressions from late- to early-successional biocrusts may alter the fraction and identity of taxa driving short-term metabolic pulses, with potential consequences for nutrient cycling dynamics in dryland soils.

## Acknowledgements

R.R. acknowledges the European Union Horizon 2020 research and innovation program under the Marie Skłodowska-Curie IF-GF Actions (MICROBIOCLIM, GA nr. 101028323), and the projects CRUST R-Forze (PID2021-127631NA-I00) and CNS2024-154916, funded by FEDER/Ministerio de Ciencia e Inovacion-Agencia Estatal de Investigacion. We thank Sasha Reed for providing access to the study sites. E.C. is supported by the USDA National Institute of Food and Agriculture and Hatch Appropriations under Project #PEN04949 and Accession #7006508. F.T.M. acknowledges support from the King Abdullah University of Science and Technology. We acknowledge the Huck Institutes’ Flow Cytometry Core Facility (RRID:SCR_024460) for the use of the BD Fortessa Flow Cytometer and MoFlo Astrios Cell Sorter for the collection of cells used for 16S rRNA analysis. We thank Rajeswaran Mani and Mitchell Koptchak for guiding us during flow cytometry and cell sorting procedures.

