## Supplementary Material for "Rainfall-induced microbial resuscitation reveals functional decoupling across biocrust succession"

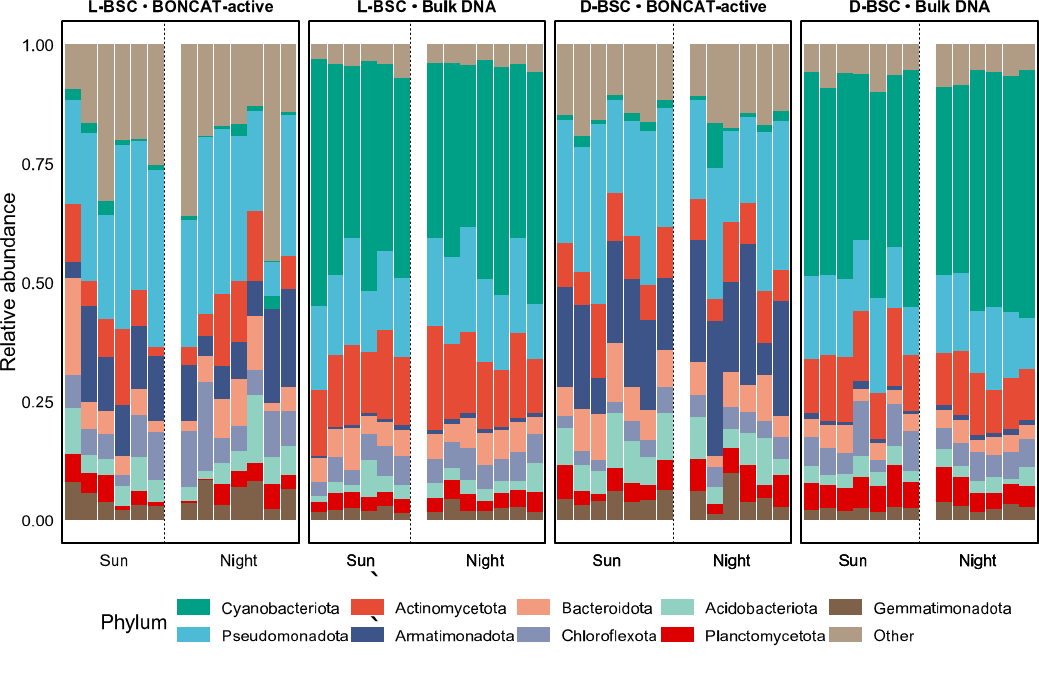

**Fig. S1**. Top 10 phyla level microbial community composition across all samples. Each stacked bar is a sample. Incubation treatment (Sun or Night) is on the X-axis. ASVs are unrarefied.

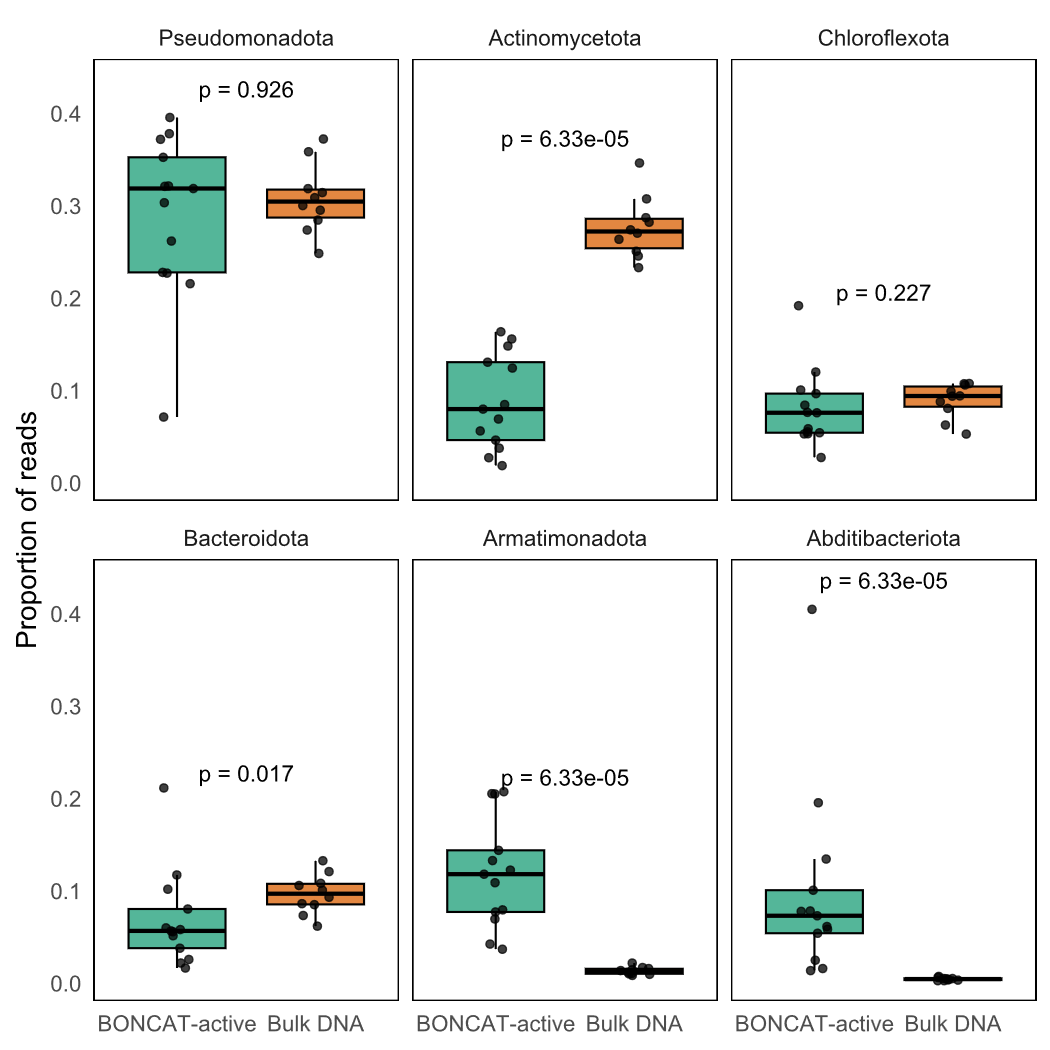

**Figure S2**. Top 6 dominant phyla in L-BSC and their enrichment in BONCAT-active vs. Bulk DNA, defined as those with the highest mean relative abundance across all samples and fractions The p-values are from Wilcoxon rank-sum tests.

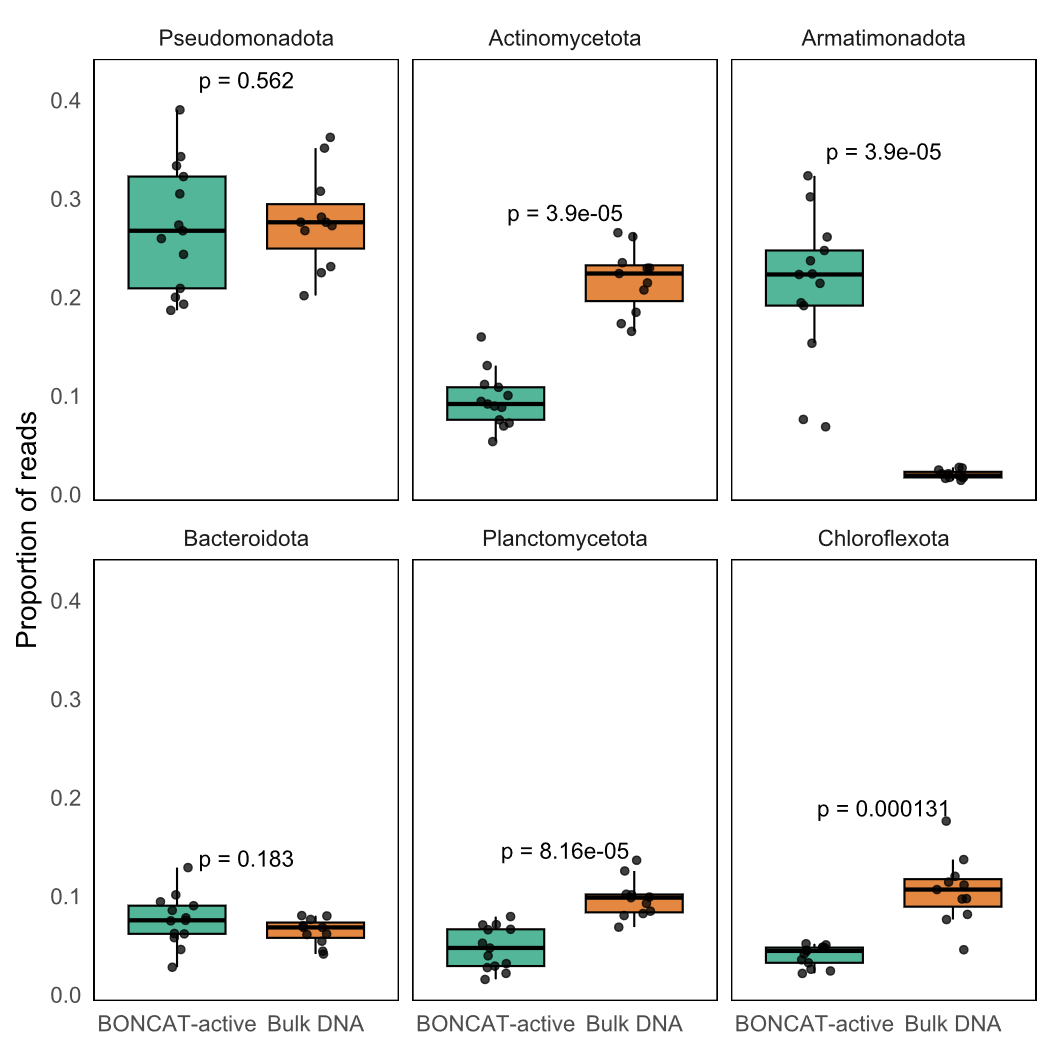

**Figure S3**. Top 6 dominant phyla in D-BSC and their enrichment in BONCAT-active vs. Bulk DNA, defined as those with the highest mean relative abundance across all samples and fractions The p-values are from Wilcoxon rank-sum tests.

**Table S1**. ANOVA results testing the effects of biocrust type and incubation conditions, and their interaction on total microbial cells. Asterisks (*) indicate p-values where factors significantly (*P* < 0.05) impacted microbial cells.

| Model: Cells_total~(BiocrustType+Incubation)^2 |  |  |
| --- | --- | --- |
| Factor | ***P*_value_** | **F** |
| BiocrustType | 1.801e-06* | 44.171 |
| IncubationConditions | 0.020* | 6.377 |
| BiocrustType:IncubationConditions | 0.021* | 6.232 |

**Table S2**. ANOVA results testing the effects of biocrust type and incubation conditions, and their interaction on BONCAT-active microbial cells. Asterisks (*) indicate p-values where factors significantly (*P* < 0.05) impacted microbial cells.

| Model: Cells_active~(BiocrustType+Incubation)^2 |  |  |
| --- | --- | --- |
| Factor | ***P*_value_** | **F** |
| BiocrustType | 9.985e-06* | 34.273 |
| IncubationConditions | 0.125 | 2.558 |
| BiocrustType:IncubationConditions | 0.545 | 0.377 |
